# Cyanobacterial NOS expression improves nitrogen use efficiency, nitrogen-deficiency tolerance and yield in Arabidopsis

**DOI:** 10.1101/2020.10.08.331702

**Authors:** Del Castello Fiorella, Foresi Noelia, Nejamkin Andrés, Lindermayr Christian, Buegger Franz, Lamattina Lorenzo, Correa-Aragunde Natalia

## Abstract

Developing strategies to improve nitrogen (N) use efficiency (NUE) in plants is a challenge to reduce environmental problems linked to over-fertilization. The nitric oxide synthase (NOS) enzyme from the cyanobacteria *Synechococcus* PCC 7335 (SyNOS) has been recently identified and characterized. SyNOS catalyzes the conversion of arginine to citrulline and nitric oxide (NO), and then 70% of the produced NO is rapidly oxidized to nitrate by an unusual globin domain in its 5'-terminus. In this study, we assessed whether SyNOS expression in plants affects N metabolism improving NUE and yield. Our results showed that transgenic Arabidopsis plants had higher primary shoot length and shoot branching when grown in N-deficient conditions and higher seed production in N-sufficient and -deficient conditions. Moreover, transgenic plants showed significantly increased NUE in both N conditions. No differences were observed in N uptake for SyNOS lines. However, SyNOS lines presented an increase in N assimilation/remobilization under low N conditions. In addition, SyNOS lines had greater N-deficiency tolerance compared to wt plants. Our results support that SyNOS expression generates a positive effect on N metabolism and seed production in Arabidopsis, and it might be envisaged as a strategy to improve productivity in crops under adverse N environments.

## INTRODUCTION

Nitrogen (N) is an important macronutrient for plants and a significant factor that limits plant growth and productivity. The great demand for increasing agricultural food production has been associated with over-application of N fertilizers to obtain better yields. Besides being expensive, the excessive use of fertilizers has large detrimental effects on the environment, biodiversity and human health. The most relevant problem is nitrate (NO_3_^−^) infiltration into groundwater by leaching, which causes eutrophication of aquatic ecosystems (Savci, 2012; Breitburg and Grégoire, 2018). It also affects soil quality and fertility, and increases greenhouse gases emissions that contribute to global warming and ozone layer depletion (Ayoub, 1999; Follett *et al*., 2010). The N use efficiency (NUE) in plants is defined as the plant seed yield relative to the amount of applied N (Xu *et al*., 2012). Improving plant NUE is crucial to increase crop yields and to reduce environmental problems linked to over-fertilization.

The study of the molecular bases that regulate plant NUE has been a main objective in plant biology research. NUE depends on both N availability in soils and the plants ability to use it efficiently. Plant NUE is composed of N uptake efficiency and N utilization efficiency, and the latter comprises both N assimilation efficiency and N remobilization efficiency (Masclaux-Daubresse *et al*., 2010). Many critical genes involved in different N metabolism steps have been manipulated to improve NUE in diverse plant species. Approaches to increase plant NUE have focused mainly on N uptake and transport by overexpression of NO_3_^−^ transporters (NTRs), and on N assimilation principally by overexpression of NO_3_^−^ reductase (NR) and glutamine synthetase (GS) genes (Good *et al*., 2004; Masclaux-Daubresse *et al*., 2010). However, efforts to increase plant productivity by augmenting N uptake or assimilation have had limited success. So far, few attempts to increase N recycling and remobilization in plants have been reported. During the reproductive stage, nutrients are remobilized and transported from source to sink organs, such as flowers, fruits and seeds. Thus, N remobilization from organic N storage strongly influences the amount of N allocated to seed production in the plant. Positive results on plant yield were achieved affecting N remobilization by overexpression of the amino acid transporter APP1 in pea and autophagy gene ATG8 in Arabidopsis (Perchlik and Tegeder, 2017; Chen *et al*., 2019). Exploring novel tools to improve NRE in plants might contribute to obtain enhanced productivity without a high demand for N supplementation.

Nitric oxide (NO) has been linked to N metabolism in plants as a signal molecule or as a source in N metabolism upon its oxidation to NO_3_^−^ by non-symbiotic hemoglobins (Dordas *et al*., 2003). In animals, NO synthase (NOS) enzymes catalyze NO and L-citrulline biosynthesis from the substrate L-arginine (Arg). NOS acts as a homodimer, and each monomer comprises two fused functional domains: an oxygenase domain (which contains a heme center and Arg and tetrahydrobiopterin (H_4_B) binding sites) at the N-terminus and a reductase domain (which contains binding sites for NADPH, FAD, and FMN) at the C-terminus. Both domains are connected by a calmodulin binding site (Griffith and Stuehr, 1995). Lately, NOS enzymes have been identified in photosynthetic microorganisms such as green algae, diatoms and cyanobacteria (Foresi *et al*., 2010; Di Dato *et al*., 2015; Kumar *et al*., 2015; Jeandroz *et al*., 2016). However, NOS sequences have not been found yet in land plant genomes (Jeandroz *et al*., 2016).

We have recently characterized the functionality of the NOS from the cyanobacteria *Synechococcus* PCC 7335 (SyNOS). SyNOS has a similar structure to animal NOS with both oxygenase and reductase domains, and contains an additional domain in the N-terminus that encodes a globin (Correa-Aragunde *et al*., 2018). SyNOS activity *in vitro* assays demonstrated that the globin domain acts as a NO dioxygenase, oxidizing NO to NO_3_^−^ (Picciano and Crane, 2019). As a result, SyNOS is able to produce NO from Arg and, in addition, to catalyze the conversion of NO to NO_3_^−^ with a release rate higher for NO_3_^−^ than for NO (Picciano and Crane, 2019). Recombinant SyNOS expression in *Escherichia coli* enhances bacterial growth rate under N-deficient conditions. Additionally, bacteria expressing SyNOS are able to grow in media containing Arg as the only N source, which indicates a possible participation of SyNOS-mediated NO/NO_3_^−^ production in N metabolism and/or assimilation (Correa-Aragunde *et al*., 2018).

In this work, we evaluated SyNOS expression as a possible strategy to increase NUE and seed yield in plants. Our working hypothesis was that SyNOS expression in plants might remobilize N from Arg storage pools, making N more available and hence favoring plant growth and development. We generated SyNOS transgenic Arabidopsis plants and analyzed their growth and seed production under high and low N conditions. Results indicate that SyNOS expression in Arabidopsis increases NUE and seed production both in N-sufficient and -deficient conditions. Furthermore, SyNOS lines have higher tolerance to N-deficiency stress compared to wt plants.

## MATERIALS AND METHODS

### Plant material and growth conditions

*Synechococcus* PCC 7335 (SyNOS) NOS nucleotide sequence was cloned in the destination plasmid pUB-DEST (Grefen *et al*., 2010), under the regulation of UBQ10 promoter. The construct was used to transform *Agrobacterium tumefaciens* strain GV3101::pMP90, and introduced into Arabidopsis plants ecotype Columbia Col-0 *rdr-6* (ABRC stock name CS24285; Luo & Chen, 2007), as previously described (Clough and Bent, 1998). The harvested seeds were stratified at 4°C for 3 days in darkness, and then cultured in ½ MS medium (Murashige and Skoog, 1962), containing 1% (w/v) agar with 1 μM ammonium glufosinate (BASTA) (Sigma) antibiotic. SyNOS homozygous lines were analyzed and selected based on BASTA resistance.

Plants were grown under long-day conditions (16/8 hours, light/dark), at 25°C, 150 μE m^−2^ s^−1^ light intensity and 60% humidity. To analyze SyNOS gene and transcript, and NOS activity, wt and SyNOS transgenic homozygous seeds were stratified and grown in 1% (w/v) agar with ½ MS medium plus 1% (w/v) sucrose for 15 days. For the experiments under different N conditions, wt plants and SyNOS homozygous lines were stratified for 3 days and sown in 0.25 l pots, one plant per pot, containing substrate with soil:perlite:vermiculite (1:1:1 v/v). N content in the soil substrate was 0.174 % (w/w). Each pot was irrigated weekly with 10 ml 4.5 mM Ca(NO_3_)_2_ (high N condition) or water (low N condition). For chlorophyll and anthocyanin quantification, the proceedings were the same except that four plants were grown per pot.

### PCR analysis

For DNA genomic extraction, 1-2 leaves from each plant were homogenized in 300 μl extraction buffer (100 mM Tris-HCl pH 8, 1.4 mM NaCl, 20 mM EDTA pH 8, 2% (w/v) CTAB and 1% (w/v) polyvinylpyrrolidone (PVP)), and then heated at 65°C for 5 min. Chloroform (150 μl) was added and centrifuged at 10,000 rpm for 2 min. Then, 180 μl isopropanol was added to the aqueous phase and the samples were incubated at room temperature for 30 min. Samples were centrifuged at 10,000 rpm for 2 min and the pellets were washed with 80% (v/v) ethanol. Genomic DNA was resuspended in sterile water. The primer sequences for SyNOS and Actin that amplify 4451 bp and 651 bp fragments respectively, are shown in Supplementary Table S1. PCR reactions were performed using 1 μl cDNA, 10 pmol of each primer and 1 μl Taq polymerase (Invitrogen) in a 20 μl reaction volume, with annealing temperature of 55°C and 35 cycles. An aliquot of 10 μl PCR products was analyzed by electrophoresis in 1% (w/v) agarose gels.

### Quantitative PCR analysis

Total RNA was extracted using Trizol isolation reagent (Invitrogen), and treated with RQ1 RNase-free DNase (Promega). One μg of total RNA was used for first-strand cDNA synthesis with an oligo(dT) primer and M-MLV reverse transcriptase (Promega). The primer sequences are listed in Supplementary Table S1. For quantitative RT-PCR, reactions were performed on a Step-one Real-time PCR machine from Applied Biosystems (California, USA) with Fast Universal SYBR Green Master Rox (Roche) to monitor double-stranded DNA synthesis. Relative transcript levels were determined for each sample and normalized against actin transcript levels.

### NOS activity

NOS activity was determined in Arabidopsis plant extracts by monitoring the conversion of [^3^H]-Arg into [^3^H]-L-citrulline as described by Bredt & Snyder (1990). Enzymatic reactions were performed at 25°C in 50 mM Tris-HCl pH 7.4, containing 50 mM L-Arg, 1 μCi [^3^H]-Arg monohydrochloride (40–70 Ci/mmol; Perkin-Elmer), 100 μM NADPH, 10 μM FAD, 2 mM CaCl_2_, 10 μg calmodulin and 100 μM H_4_B in a volume of 40 μl. Reactions were initiated adding 20 μg total plant proteins, and stopped after 30 min with 400 μl ice-cold stop buffer (20 mM sodium acetate pH 5.5, 1 mM L-citrulline, 2 mM EDTA and 0.2 mM EGTA). Samples were applied to columns containing 1 ml Dowex AG50W-X8, Na^+^ form (100–200 mesh; Bio-Rad), pre-equilibrated with stop buffer. L-citrulline was eluted using 2 ml distilled water. Eluate aliquots (0.5 ml) were dissolved in 10 ml scintillation liquid, and radioactivity was measured using a Beckman LS 3801 liquid scintillation system.

### Determination of nitrate and protein

NO_3_^−^ and protein quantification in seeds was performed according to Lam et al. (2003). Seeds (50 mg) were ground in liquid N_2_ with the addition of glass beads, and then resuspended in 200 μl extraction buffer (250 mM NaCl and 50 mM potassium phosphate buffer pH 7). After centrifugation at 12,000 g for 5 min at 4°C, the supernatants were used for quantifications.

NO_3_^−^ content was determined as described by Cataldo et al. (1975). Samples (10μl) were incubated with 40μl salicylic acid (5mg.ml^−1^ in H_2_SO_4_) for 20min and then neutralized with 950μl 2N NaOH. Absorbance at 410nm was measured. NO_3_^−^ concentration was estimated by comparison with a NO_3_^−^ standard calibration curve. Protein content in samples was determined with the Bradford method (Bradford, 1976).

### Chlorophyll and anthocyanin contents

Chlorophyll from entire rosettes was extracted in 100% (v/v) cold ethanol (0.45 g tissue per 10 ml solution) at 4°C in darkness for 24 h. Total chlorophyll content was calculated measuring the absorbance at 470, 648 y 664 nm and according to the equations established by Lichtenthaler (1987).

To estimate anthocyanin concentration, entire rosettes were incubated in 80% (v/v) methanol containing 0.01 M HCl (0.45 g tissue per 10 ml solution) at 4°C in darkness for 24 h. Then, 300 μl of extract was added to 200 μl H_2_O and 100 μl chloroform, and centrifuged (13,000 g for 10 min). Anthocyanin content was estimated by calculating the difference between absorbance at 540 and 630 nm in the methanolic phase (Diaz *et al*., 2006).

### Determination of ^15^N content

Plants were grown in hydroponic complete ATS medium for 21 days and then transferred to high N (9 mM NO_3_^−^) or low N (0.5 mM NO_3_^−^) conditions. To analyze NO_3_^−^ uptake, in both high and low N solution, NO_3_^−^ was enriched with 20 atom% ^15^N. After switching the N conditions, atmospheric NO incorporation was analyzed in plants exposed to 90 ppbv ^15^NO. Kinetics of ^15^NO_3_^−^ uptake and atmospheric ^15^NO incorporation were determined at 3, 6 and 9 days. Plant material was dried at 60°C overnight and ground to powder using a ball mill (Tissue Lyser II, Qiagen, Venlo, Netherlands). Aliquots of about 1.5-2 mg were transferred into tin capsules (IVA Analysentechnik, Meerbusch, Germany). ^15^N abundance was determined with an Isotope Ratio Mass Spectrometer (IRMS) (delta VAdvantage, Thermo Fisher, Dreieich, Germany) coupled to an Elemental Analyzer (Euro EA, Eurovector, Milano, Italy) as described by Kuruthukulangarakoola et al. (2017).

### Total N in plants and soil samples

Total N in seed and plant samples was determined using Dumas method (McGill *et al*., 2007) by dry combustion at 950°C and thermo-conductivity measurement using a TruSpec N analyzer (LECO Corporation, St. Joseph, Michigan, USA).

The same method was used for N quantification in soil samples. NUE values were determined as the ratio between seed production (g) per plant and total N content (g) per pot provided by the substrate. Total N in soil substrate includes initial N content in the substrate plus the weekly NO_3_^−^ supplement (10 weeks).

### Statistical analysis

Statistical analyses were conducted with R software (version 3.5.1; R Foundation for Statistical Computing). The model assumptions used were tested with graphic analysis and Shapiro-Wilk test. The data sets that did not comply with normality were evaluated with Gamma distribution. The discrete variables were analyzed with Poisson distribution. Statistical significance was determined by ANOVA with post hoc Dunnett’s test for multiple comparison analyses, or Student's t test for pairwise comparisons.

## RESULTS

### Characterization of transgenic Arabidopsis plants expressing SyNOS gene

Arabidopsis plants were transformed with a DNA sequence coding SyNOS under the regulation of the ubiquitin constitutive promoter (Fig. 1A). Two SyNOS homozygous lines that represent different transformation events, Sy6 and Sy7, were selected. Figures 1B and C show the detection of the *SyNOS* transgene and transcript by PCR and real-time PCR respectively, in SyNOS transgenic lines. SyNOS transcript levels in Sy7 line were higher than those in Sy6 line (Fig. 1C). NOS activity in leaves was determined by measuring [^3^H]-citrulline production with [^3^H]-Arg as the substrate. SyNOS transgenic lines showed higher NOS activity compared to wt plants (Fig. 1D). The higher NOS activity observed in Sy7 line was in agreement with the higher SyNOS expression compared to Sy6 line. Overall, these results suggest that SyNOS is active in the transgenic plants.

**Figure 1.**
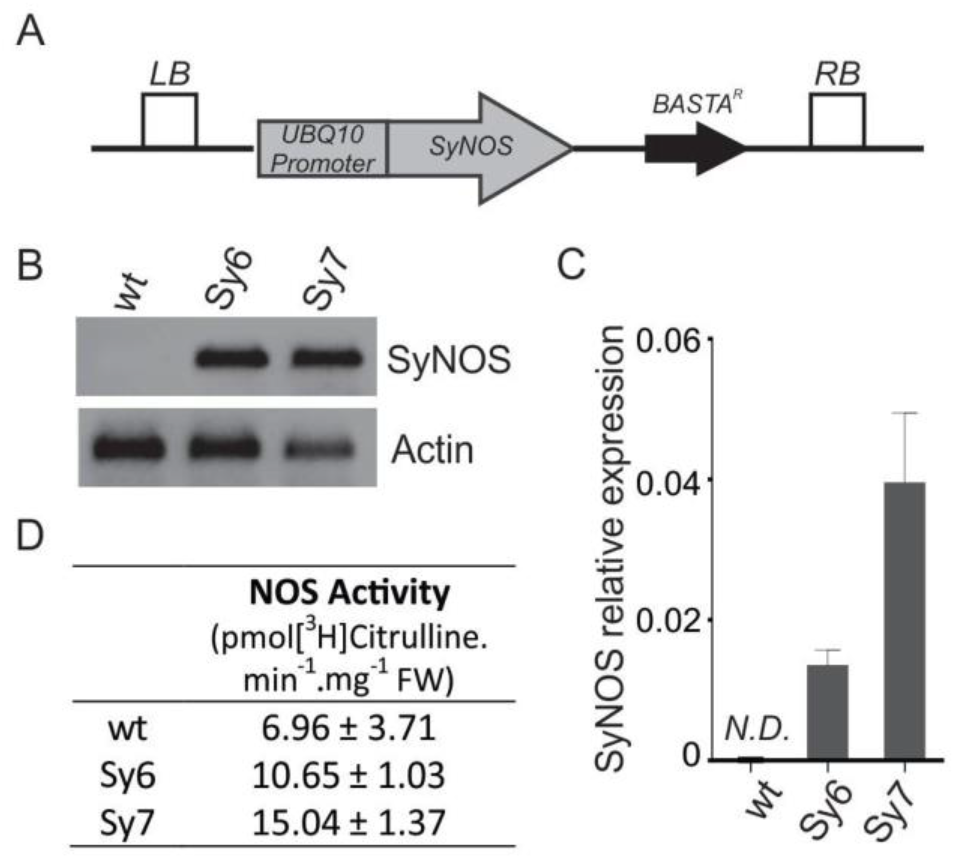
Expression analysis of NO synthase from *Synechococcus* PCC 7335 (SyNOS) in Arabidopsis. (A) Construct used to obtain transgenic Arabidopsis plants expressing SyNOS gene. UBQ10, ubiquitin promoter; SyNOS, full-length DNA encoding *SyNOS*; BASTA^R^, phosphinothricin-*N* - acetyltransferase gene that confers resistance to the herbicide BASTA; LB and RB, left and right T-DNA border sequences, respectively. (B) Analysis of the presence of *SyNOS* transgene by PCR in wt and in two independent transgenic lines (Sy6 and Sy7). Actin expression was evaluated as a loading control. (C) *SyNOS* transcript levels analyzed by qPCR. Transcript levels are normalized using actin as reference gene. N.D.: no detection. (D) NOS activity in leaf extracts from wt and transgenic lines determined by measuring [^3^H]-Citrulline production from [^3^H]-Arg. FW, fresh weight. Values are mean ± SE from three independent experiments using 5-plants rosette pool (n=3).

### SyNOS transgenic lines have higher seed production and NUE in both high and low N conditions

In order to analyze the transgenic lines development and productivity, plants were grown under two different N availability conditions. Seeds were sown in pots containing substrate with soil:perlite:vermiculite (1:1:1) and weekly supplemented with NO_3_^−^ (high N condition) or only irrigated with water (low N condition). Figure 2 (A-C) shows that N deficiency caused a decrease in primary shoot length, number of secondary shoots and seed production in wt plants, as previously reported (Martin *et al*., 2002; Lemaître *et al*., 2008; North *et al*., 2009; Jong *et al*., 2014). On the other hand, under low N conditions, SyNOS transgenic lines showed increased primary shoot length, shoot branching and seed production per plant with respect to wt (Fig. 2A, B and C). Under high N conditions, a statistically significant increase was observed in shoot branching for Sy7 line, the transgenic line with higher NOS activity detected (Fig. 2A and B). Furthermore, both SyNOS lines had higher seed production compared to wt plants under high N conditions (Fig. 2C). Regarding vegetative development, SyNOS lines showed no statistical differences in rosette fresh weight, dry weight and water content under both N conditions compared to wt (Supplementary Fig. S1). NUE in transgenic and wt plants was calculated as the ratio between seed production and total N content in the substrate. SyNOS lines showed higher NUE values compared to wt plants in both N conditions (Table 1), which indicates that SyNOS expression affects N metabolism in transgenic plants, improving NUE and seed yield.

**Figure 2.**
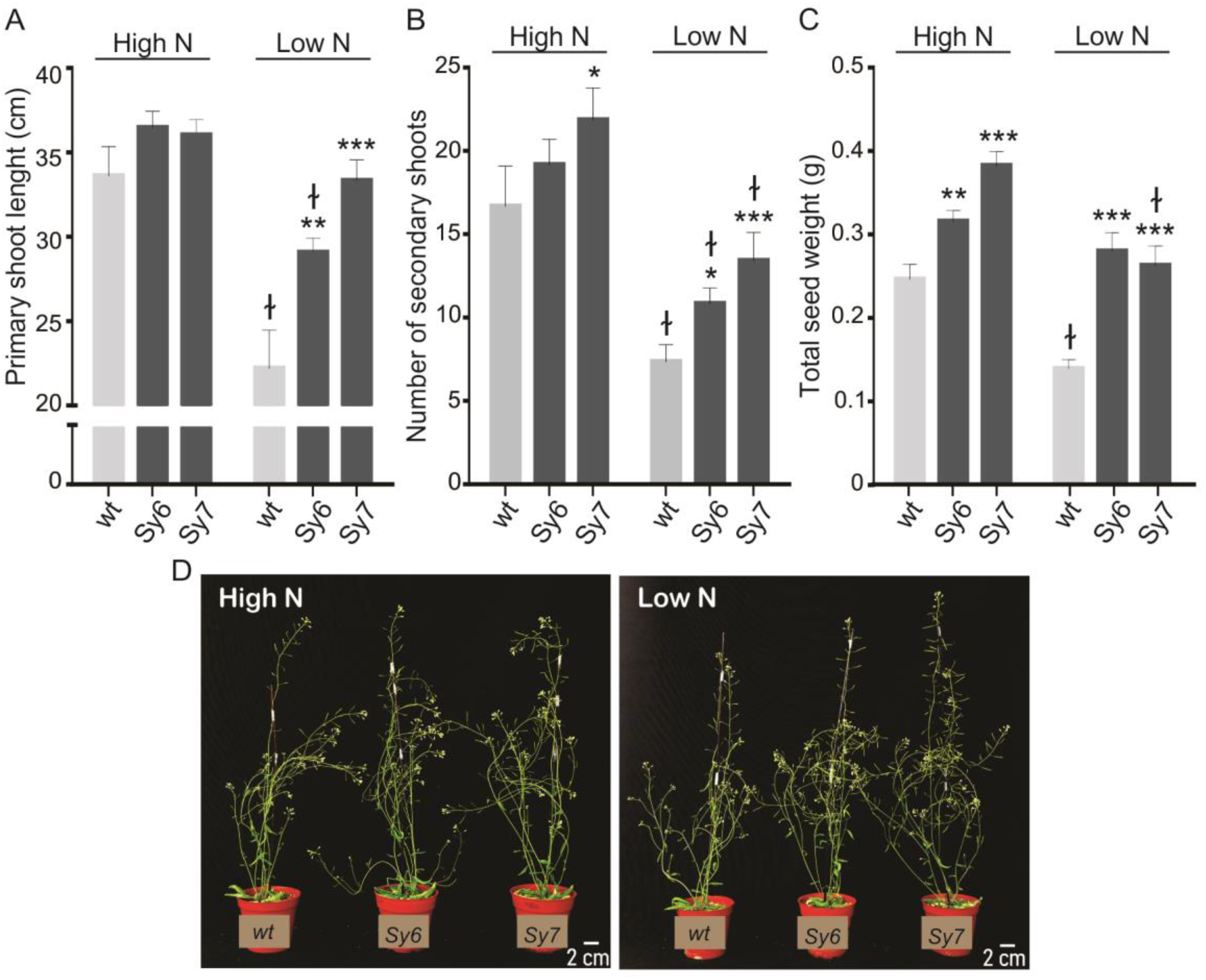
Phenotypic characterization of SyNOS transgenic Arabidopsis plants growing in different N conditions. Plants were grown in pots containing substrate with soil:perlite:vermiculite (1:1:1) and supplemented weekly with 10 ml per pot of 4.5 mM Ca(NO_3_)_2_ for high N or water for low N conditions. (A) Primary shoot length and (B) number of secondary shoots at 30 and 35 days after sowing (DAS) respectively. (C) Total seed weight per plant. (D) Picture of plants growing 40 DAS in high N and low N conditions. Values are means ± SE (n=6, for each parameter). Three independent experiments (n=6 each) were performed with similar results. Asterisks indicate significant differences compared to wt grown in the same N conditions (ANOVA, Dunnett’s post hoc test, *p<0.1, **p<0.05, ***p<0.01). ƚ indicates significant differences between the different N conditions in each line (Student’s t-test, p<0.05).

**Table 1:** N use efficiency (NUE) of plants growing in high and low N conditions.

| Genotype | NUE |  |
| --- | --- | --- |
|  | High N | Low N |
| **wt** | 0.7 ± 0.05 | 1.21 ± 0.08 |
| **Sy6** | 1.4 ± 0.11 ** | 1.55 ± 0.06 * |
| **Sy7** | 1.3 ± 0.12 ** | 1.88 ± 0.08 ** |

NUEs were calculated as the ratio between seed production (mg) per plant and total N content (mg) per pot provided by the substrate. Values are means ± SE (n=6). Asterisks indicate significant differences compared to wt for the same condition (ANOVA, Dunnett's post hoc test, \*p<0.05, \*\*p<0.01).

Besides yield, the percentages of total N and protein in grains represent other important agronomic traits. Numerous studies have reported a negative correlation between yield and seed quality, showing that breeding cultivars with higher yield result in a decrease in protein and metabolite contents in seeds (Beninati and Busch, 1992; Lemaître *et al*., 2008). To check the transgenic seeds quality we analyzed the seeds harvested from plants grown in low N conditions, where greater phenotypic differences among wt and SyNOS lines had been observed. Table 2 shows that SyNOS expression did not affect seed size or total N metabolite levels. Intriguingly, Sy7, the line with higher NOS activity, presented less NO_3_^−^ but more protein content in seeds compared to wt, perhaps due to an augmented NO_3_^−^ incorporation into amino acids (Table 2). Moreover, 100% germination was observed within 72 h, indicating that SyNOS expression did not affect seed viability either. These results indicate that the increase in yield does not affect seed quality negatively in SyNOS lines.

**Table 2.** Characterization of seeds harvested from wt and SyNOS transgenic Arabidopsis plants under low N condition.

| | **Seed**<br>**Perimeter**<br>(mm) | **Nitrate**<br>($\mu\text{mol.g}^{-1}$ seed) | **Protein**<br>($\text{mg.g}^{-1}$ seed) |
| --- | --- | --- | --- |
| **wt** | $1.35 \pm 0.011$ | $0.82 \pm 0.02$ | $53.9 \pm 2.4$ |
| **Sy6** | $1.31 \pm 0.010$ | $0.75 \pm 0.03$ | $50.0 \pm 1.5$ |
| **Sy7** | $1.28 \pm 0.011$ | $0.72 \pm 0.01^*$ | $90.2 \pm 2.9^*$ |

Seed perimeters were measured using ImageJ software. Values are means and ± SE (n=60). Values of nitrate and protein are means and ± SE (n=6). These experiments were performed three times (n=6) with similar results. Asterisks indicate significant differences compared to wt plant (ANOVA with post hoc Dunnett's method, \*p<0.05).

### SyNOS plants present higher chlorophyll and lower anthocyanin levels than wt under low N conditions

Plants are able to acclimate to N restriction by developing various adaptive responses. Photosynthesis reduction and anthocyanin accumulation are two important adaptive responses to face N deficiency (Diaz *et al*., 2006). The reduced photosynthetic capacity is correlated with an increased chlorophyll degradation by senescence processes (Thomas and de Villiers, 1996). On the other hand, anthocyanin plays a protective role by scavenging reactive oxygen species (ROS) in senescent leaves susceptible to light damage (Lea *et al*., 2007). To assess the tolerance to N deficiency in transgenic and wt plants, chlorophyll and anthocyanin pigments were measured in leaves from plants growing under low N conditions (Fig. 3). In accordance with NOS activity in the transgenic lines, Sy7 line displayed 38% more chlorophyll content than wt, while Sy6 line showed an increase of only 22% (Fig. 3A). In addition, anthocyanin levels were 43% and 67% lower in Sy6 and Sy7 respectively compared to wt (Fig. 3B). These results suggest that SyNOS lines have or perceive a better N status than wt plants.

**Figure 3.**
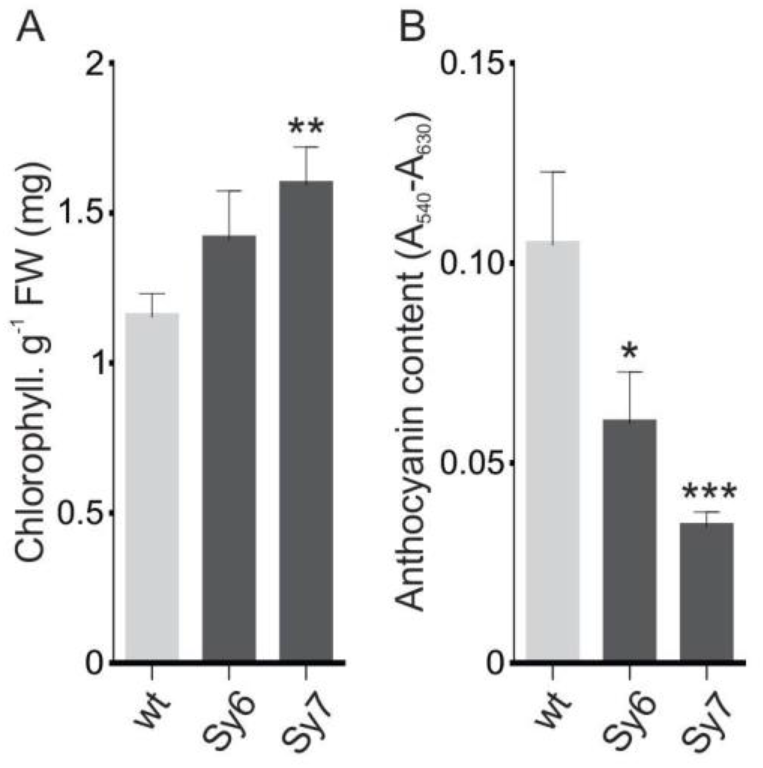
Chlorophyll and anthocyanin contents in wt and SyNOS transgenic lines growing in low N conditions. (A) Chlorophyll and (B) anthocyanin contents were quantified in leaves from plants at 30 DAS in low N conditions (four plants per pot). FW, fresh weight. Values are means ± SE (n=5). Asterisks indicate significant differences compared to wt (ANOVA, Dunnett’s post hoc test, *p<0.1, *p<0.05, **p<0.01).

### SyNOS heterologous expression improves plant N remobilization

To further investigate how SyNOS expression improves NUE in plants, we analyzed different processes involved in the use of N in wt and transgenic lines. First, NO_3_^−^ uptake and atmospheric NO incorporation were determined in wt and SyNOS plants (Fig. 4). Plants were grown for 3 weeks in a hydroponic culture containing N and then transferred to two N availability conditions: high N (9 mM NO_3_^−^) or low N (0.5 mM NO_3_^−^). To analyze NO_3_^−^ uptake, NO_3_^−^ was enriched with 20 atom% ^15^N in both high and low N solutions. To evaluate NO-fixation, plants were exposed to 90 ppbv ^15^NO after switching the N conditions. Figure 4 shows that SyNOS transgenic and wt plants had similar NO_3_^−^ uptake and atmospheric NO incorporation rates under both N conditions. The kinetics of ^15^N incorporation of NO_3_^−^ and NO are shown in Supplementary Fig. S2. These results indicate that the increase in growth and seed production observed in SyNOS lines is not due to a higher N uptake of NO_3_^−^ from soil or atmospheric NO. Interestingly, all lines had greater NO incorporation under low N conditions with respect to high N conditions, suggesting that plants raise atmospheric NO fixation when N is scarce in the soil (Fig. 4B).

**Figure 4.**
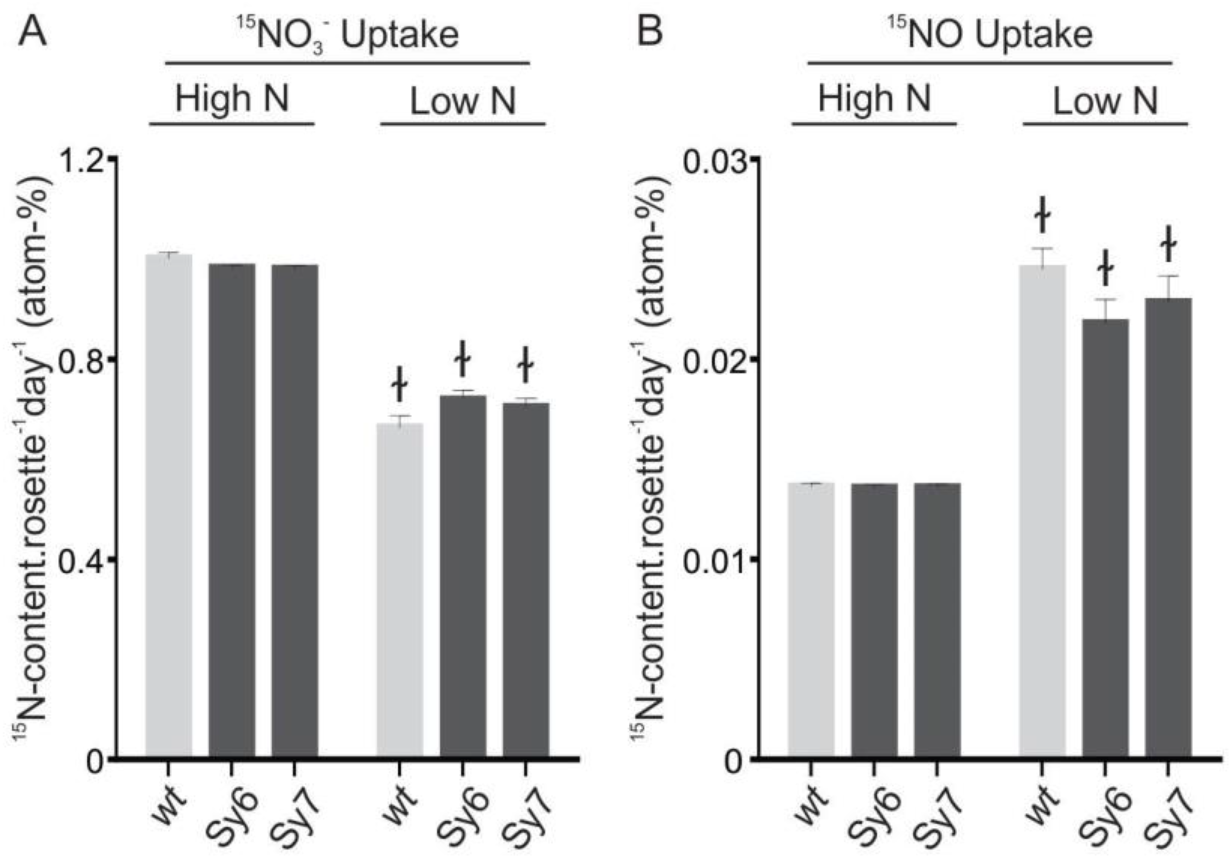
^15^NO_3_^−^uptake and atmospheric ^15^NO incorporation in SyNOS transgenic plants. Plants were grown in complete ATS hydroponic culture for 21 days and then transferred to high N conditions (9 mM NO_3_^−^) or low N conditions (0.5 mM NO_3_^−^). (A) To analyze NO_3_^−^ uptake, N solution was enriched with 20 atom% ^15^NO_3_^−^ in both high and low N conditions. (B) After switching the N conditions, atmospheric NO incorporation was analyzed in plants exposed to 90 ppbv ^15^NO. The incorporation of ^15^N per day was calculated based on the ^15^N data 9 days after changing the N solutions. Values are means ± SE (n=6). ƚ indicates significant differences between the different N conditions in each line (Student’s t-test, p<0.05).

Once NO_3_^−^ is internalized, it is used to synthesize organic macromolecules. Primary paths in N assimilation involve NO_3_^−^ reduction to nitrite and ammonium by NR and nitrite reductase (NiR) enzymes, followed by ammonium assimilation into amino acids by the GS / glutamate synthase (GOGAT) cycle (Masclaux-Daubresse *et al*., 2010). To evaluate N assimilation, the above-mentioned transcripts were evaluated by qPCR in leaves from wt and SyNOS plants growing in low N conditions. Figure 5 shows that Sy7 line presented statistically higher transcript levels of NR, NiR, GS and GOGAT compared to wt plants; while Sy6 line showed a slight increase in all the analyzed transcript levels with respect to wt (Fig. 5), in agreement with lower SyNOS transcript levels and activity (Fig. 1C and D). These results indicate that higher SyNOS expression in plants affects, and possibly increases, N assimilation.

**Figure 5.**
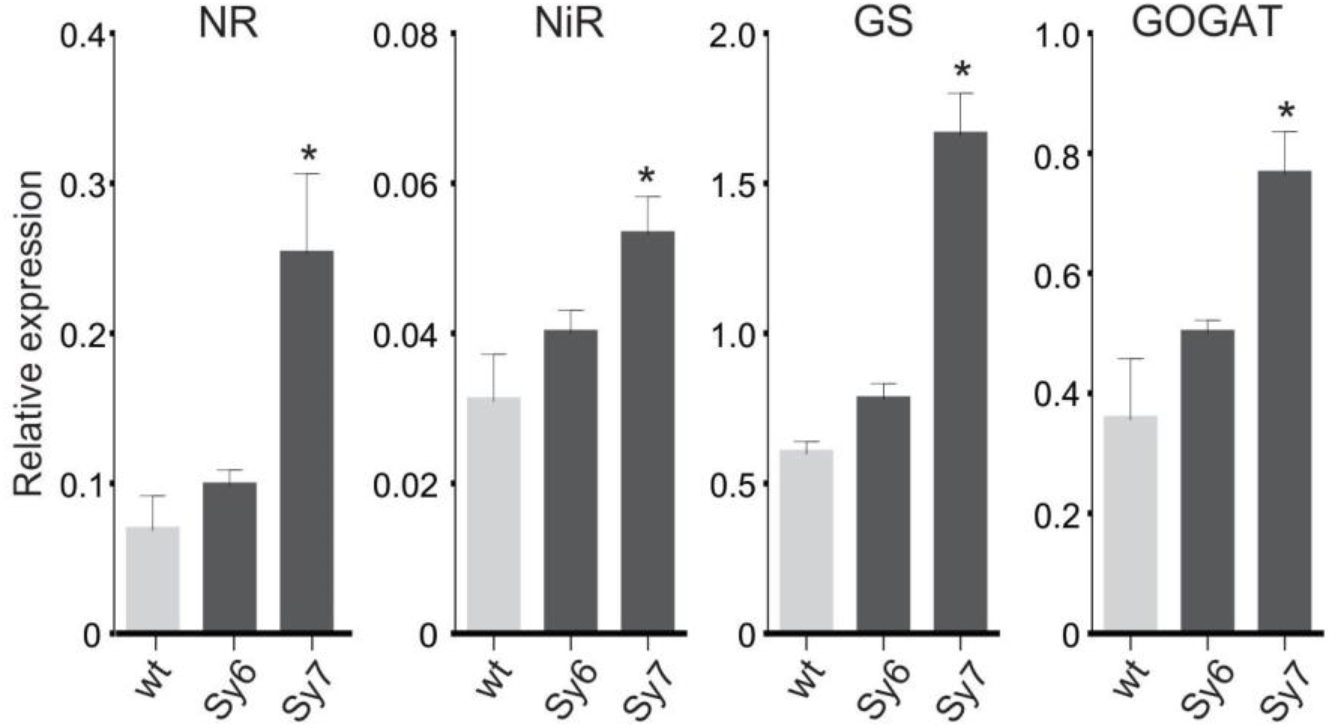
Analysis of transcript levels of genes involved in N assimilation in wt and SyNOS plants growing in low N conditions. The transcript levels of *NR* (*NiA1*), *NiR, GS2* and *GOGAT* were analyzed by qPCR in leaves from plants growing for 30 DAS in low N conditions. Transcript levels are normalized using actin as reference gene. Values are means ± SE (n=4). Asterisks indicate significant differences compared to wt (ANOVA, Dunnett’s post hoc test, *p<0.05).

Finally, we analyzed N remobilization during seed filling in plants growing in low N conditions (Fig. 6). Total N was quantified in total seeds and vegetative remains after harvest. Total N content in whole plants was similar among lines (Fig. 6A). This result was expected since no difference was detected in N uptake. On the other hand, statistical differences were observed when N partitioning to different organs in wt and SyNOS plants was analyzed (Fig. 6B). N content in SyNOS plants was lower in vegetative remains and higher in total seeds compared to wt plants. The differences were statistically significant for Sy7 line and to a lesser extent in Sy6 line (Fig. 6). These results show that N remobilization to seed filling is more efficient in SyNOS lines than wt plants and its effect is greater in the SyNOS line with higher NOS activity.

**Figure 6.**
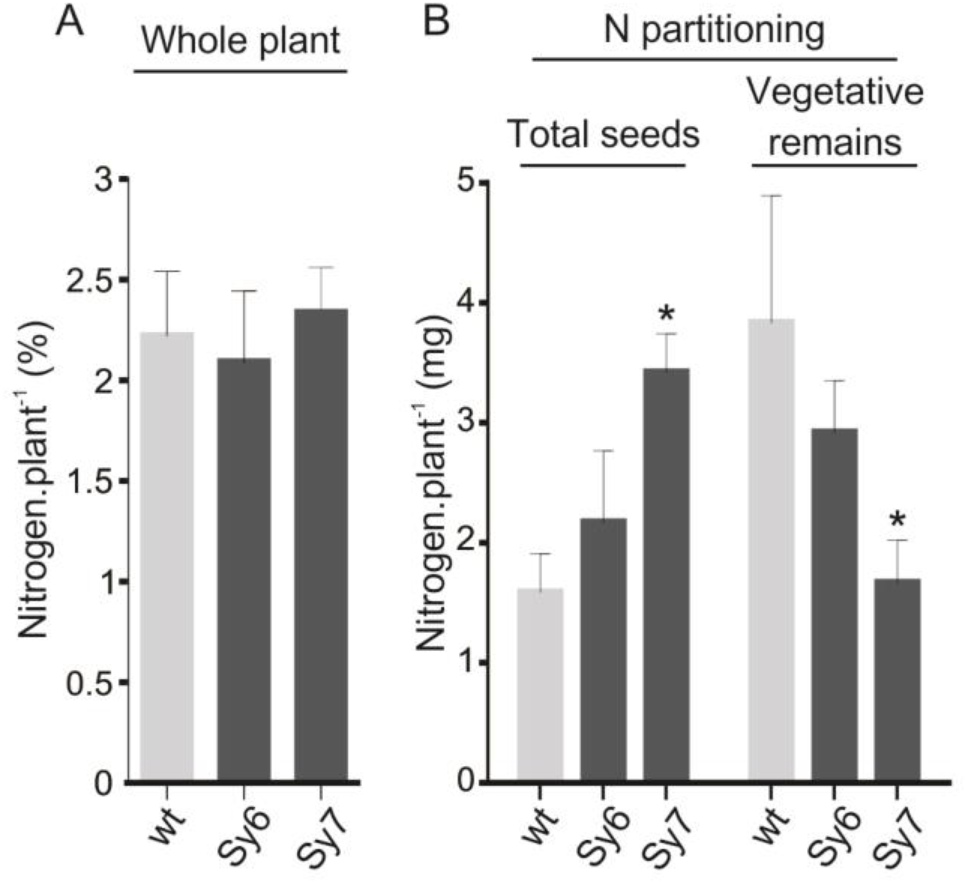
N partitioning during seed filling in SyNOS transgenic lines and wt plants growing in low N conditions. N contents in (A) the whole plant and (B) in total seeds and vegetative remains from plants growing in N-deficient conditions were quantified with the Dumas method (McGill *et al*., 2007). Values are means ± SE (n=6). Asterisks indicate significant differences compared to wt (ANOVA, Dunnett’s post hoc test, *p<0.05).

## DISCUSSION

Nitrogen (N) is a critical nutrient for plant growth and development and it is extensively used to maximize crop yields. Arg is considered an important organic N storage and transport form in plants due to its high nitrogen to carbon (N/C) ratio. Arg catabolism by arginase enzymes in plants provides urea and ornithine, and then urea is broken down into ammonium and CO_2_ by ureases. Ammonium is finally assimilated into glutamate while CO_2_ may be used in photosynthesis. Thus, N provided by Arg catabolism can be used to meet the developing organs metabolic demands (Winter *et al*., 2015). The contribution of Arg-derived N in vegetative development and seed production has been demonstrated by characterization of plant mutants defective in arginase activity (Ma *et al*., 2013; Liu *et al*., 2018). NOS from *Synechococcus* PCC 7335 (SyNOS) is a genuine Arg-degrading enzyme that produces citrulline and NO_3_^−^/NO (Correa-Aragunde *et al*., 2018; Picciano and Crane, 2019). We propose that SyNOS expression may remobilize N from Arg internal storage pools promoting NUE, plant growth and productivity in Arabidopsis. In this work, it has been shown that SyNOS expression in Arabidopsis results in an increased reproductive tissues growth, seed production and NUE compared to wt growing in different N conditions. Additionally, seed quality was not adversely affected in the transgenic lines. Interestingly, the positive effect of SyNOS expression on seed production and NUE was more significant under low N conditions. In addition, SyNOS expression conferred tolerance to N-deficiency stress in plants preventing chlorophyll degradation and diminishing anthocyanin production under low N conditions. These results suggest that SyNOS expression could be an effective strategy for improving NUE and yield in plants with low exogenous N application.

Promising results in plant productivity have been reported by the manipulation of Arg catabolism. Ma et al., (2013) showed that arginase (OsARG) overexpression in rice increases grain number under N limited conditions, showing an important participation of Arg catabolism in N recycling. According to this report, OsARG expression in cotton enhances the quality and length of cotton fibers, an important agronomic trait in this crop (Meng *et al*., 2015). These works are of special interest since they show that manipulating downstream Arg-dependent N-remobilization steps may be key to obtain better plant yields both in monocots and in dicots. Our results show that wt and SyNOS lines have similar rates in NO_3_^−^ and atmospheric NO uptake under high and low N conditions. N content in the whole plant remains invariable throughout the life cycle grown in low N conditions. In addition, SyNOS lines remobilized more N to the seed filling compared to wt plants, with significant differences in the SyNOS line with higher NOS activity. These results indicate that SyNOS expression in Arabidopsis stimulates N remobilization increasing seed production. Additional studies have shown positive results in plant yield for canola and rice by increasing Alanine biosynthesis, another N storage amino acid in plants (Good *et al*., 2007; Shrawat *et al*., 2008). In summary, we propose that SyNOS activity might act as an additional Arg catabolic pathway enhancing N recycling in plants. Thus, SyNOS expression in plants may enhance the efficiency of global N and increase seed yield even in low N conditions. Thereby, it could contribute to economize N in plants and decrease exogenous N demand.

Additionally, the line with higher SyNOS expression (Sy7) showed increased levels of the N assimilation transcripts NR, NiR, GS2 and GOGAT under low N conditions. Thus, it could be suggested that N provided from Arg catabolism in the transgenic lines might be used for macromolecule biosynthesis in plants, although further investigation is required. Besides the essential role in plant primary metabolism, NO_3_^−^ has been proposed as an important signal molecule in various processes throughout the plant life (Wang *et al*., 2000, 2003, 2004; Scheible *et al*., 2004; Krapp *et al*., 2011). SyNOS-derived NO_3_^−^ may also be regulating N response processes in plants, improving the adaptation to N limitation or perceiving a better N status. SyNOS activity also produces citrulline and NO. Citrulline has a high N/C ratio and has been considered a key molecule in N recycling (Joshi and Fernie, 2017). Several amino acid transporters have affinity for citrulline (Fischer *et al*., 2002) indicating its participation in N transport throughout the plant. Citrulline produced by NOS activity in transgenic plants could also take part in the regulation of N recycling and transport, mainly when protein turnover and amino acid recycling are augmented.

NO is a signal molecule that participates in various development and stress tolerance processes in plants (Del Castello *et al*., 2019; Kolbert *et al*., 2019). NO has also been proposed as a signal messenger in tolerance to N-deficiency stress by preserving photosynthetic pigments (Jasid *et al*., 2009; Kováčik *et al*., 2009). Furthermore, some studies have shown that NO could be a potential source of N in plants, since it can be oxidized to NO_3_^−^ by phytoglobins. Overexpression of non-symbiotic haemoglobin 1 and 2 (GLB1 and GLB2) in Arabidopsis and barley allows channeling of atmospheric NO into N metabolism improving plant growth (Kuruthukulangarakoola *et al*., 2017; Zhang *et al*., 2019). Conversely, SyNOS expression does not affect NO incorporation rate in Arabidopsis, suggesting that SyNOS globin domain would not fix external NO in the transgenic plants. Recently, Nejamkin et al. (2020) has shown that heterologous expression of NOS from *Ostreococcus tauri* (OtNOS) in tobacco increases growth rate, number of flowers and seed yield, although only under N-sufficient conditions. Unlike tobacco expressing OtNOS, which depends on N supplementation to exhibit a positive phenotypic effect, SyNOS Arabidopsis lines presented increased yield even under N-deficient conditions. OtNOS is an enzyme with an ultrafast NO production (Weisslocker-Schaetzel *et al*., 2017) thus, plants expressing OtNOS could be wasting N as gaseous NO. Conversion of NO to NO_3_^−^ by SyNOS globin domain could allow a more efficient incorporation and recycling of internal N. Furthermore, according to *in vitro* assays, SyNOS releases approximately 25% of NO (Picciano and Crane, 2019). Low levels of SyNOS-derived NO in transgenic lines could act as antioxidants to protect photosynthetic pigments from degradation during plant development under low N availability conditions.

## CONCLUSION

SyNOS expression in plants might allow N remobilization from organic N storages, providing more NO_3_^−^ availability, which in turn can act as a N source or signal molecule improving the plant nutrient status and seed production. Furthermore, SyNOS activity generates NO that could positively affect diverse physiological processes during plant development or stress tolerance, and citrulline that could also maximize N recycling and transport. Interestingly, SyNOS plants have higher NUE in N-sufficient and -deficient conditions, resulting in increased seed production. Particularly, under N-limited conditions, SyNOS plants show enhanced N assimilation and remobilization. In addition, SyNOS lines have higher tolerance to N-deficiency stress than wt plants. Overall, SyNOS expression in plants seems to be an interesting strategy to improve crops and reduce contamination caused by the excessive use of N fertilizers.

## ABBREVIATIONS

Arg: Arginine
GOGAT: Glutamate synthase
GS: Glutamine synthetase
NO_3_^−^: Nitrate
NR: Nitrate reductase
NiR: Nitrite reductase
NO: Nitric oxide
NOS: Nitric oxide synthase
SyNOS: Nitric oxide synthase from *Synechococcus* PCC 7335
OtNOS: Nitric oxide synthase from *Ostreococcus tauri*
N: Nitrogen
NUE: Nitrogen use efficiency

## SUPPLEMENTARY DATA

**Fig. S1:** Fresh weight, dry weight and water content in rosette leaves from wt plants and SyNOS lines grown in different N conditions

**Fig. S2:** Kinetics of ^15^NO_3_^−^ and atmospheric ^15^NO uptake in SyNOS-transgenic plants

**Table S1:** Primers used for PCR analysis

## DATA AVAILABILITY STATEMENT

All data supporting the findings of this study are available within the paper and within its supplementary materials published online.

## ACKNOWLEDGEMENTS

This research was supported by the Agencia Nacional de Promoción Científica y Tecnológica (ANPCyT, 2927/2015 to L.L.), the Consejo Nacional de Investigaciones Científicas y Técnicas (CONICET, PIP 0646/2015 to N.C.-A.) and institutional grants from the Universidad Nacional de Mar del Plata (UNMdP), Argentina. F.D.C. acknowledges the support of the German Academic Exchange Service (DAAD)-short term Fellowship. L.L., N.C.A. and N.F. are members of the research staff and F.D.C. and A.N. are PhD fellows from CONICET, Argentina.

## AUTHOR CONTRIBUTIONS

L.L. and C.A.N. conceived the original screening and research plan. C.A.N. and F.N. supervised the experiments. D.C.F performed the experiments. D.C.F. and C.A.N. designed the experiments and analyzed the data. N.A. contributed to statistical analysis and data analysis. B.F. performed the 15N experiments. L.C. supervised the 15N experiments and contributed to data analysis. D.C.F. conceived the project and wrote the article with contributions of all the authors. C.A.N. supervised and completed the writing. C.A.N. agrees to serve as the author responsible for contact and ensures communication.

