## Supplementary Data for "Cyanobacterial NOS expression improves nitrogen use efficiency, nitrogen-deficiency tolerance and yield in Arabidopsis"

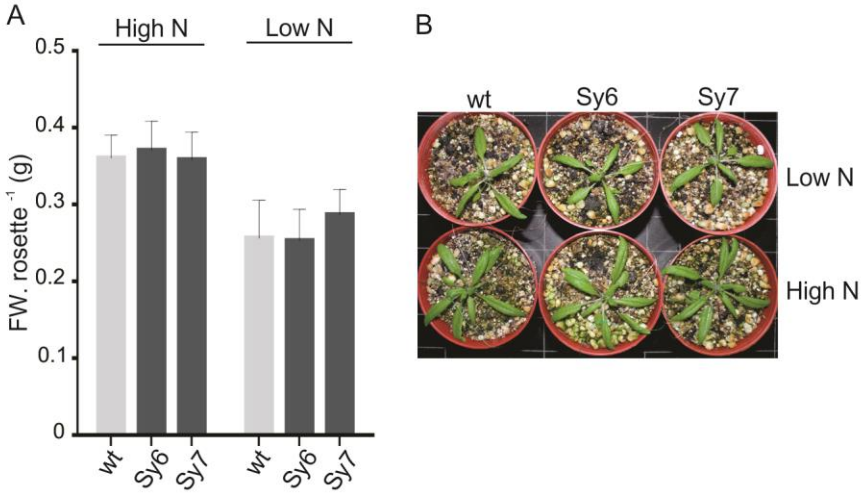


C

| Genotype | High N | | Low N | |
|---|---|---|---|---|
| | % DW | % Water | % DW | % Water |
| wt | 8 ± 0.2 | 91 ± 0.2 | 9 ± 0.9 | 90 ± 0.9 |
| Sy6 | 7 ± 0.08 | 92 ± 0.08 | 8 ± 0.4 | 91 ± 0.4 |
| Sy7 | 8 ± 0.1 | 91 ± 0.1 | 8 ± 0.1 | 91 ± 0.1 |

**Figure S1: Fresh weight, dry weight and water content in rosette leaves from wt plants and SyNOS lines grown in different N conditions**

Plants were grown in pots containing soil:perlite:vermiculite (1:1:1) and supplemented weekly with 4.5 mM $Ca(NO_3^-)_2$ (high N condition) or water (low N condition). (A) Fresh weight of rosette plants growing in high and low N conditions. (B) Picture of plants grown for 20 DAS. (C) Whole plant rosettes were dried at 45°C for 4 hours and then the dry weight (DW) was measured. Water content was determined by calculating the difference between FW and DW. Values are means ± SE (n=6).

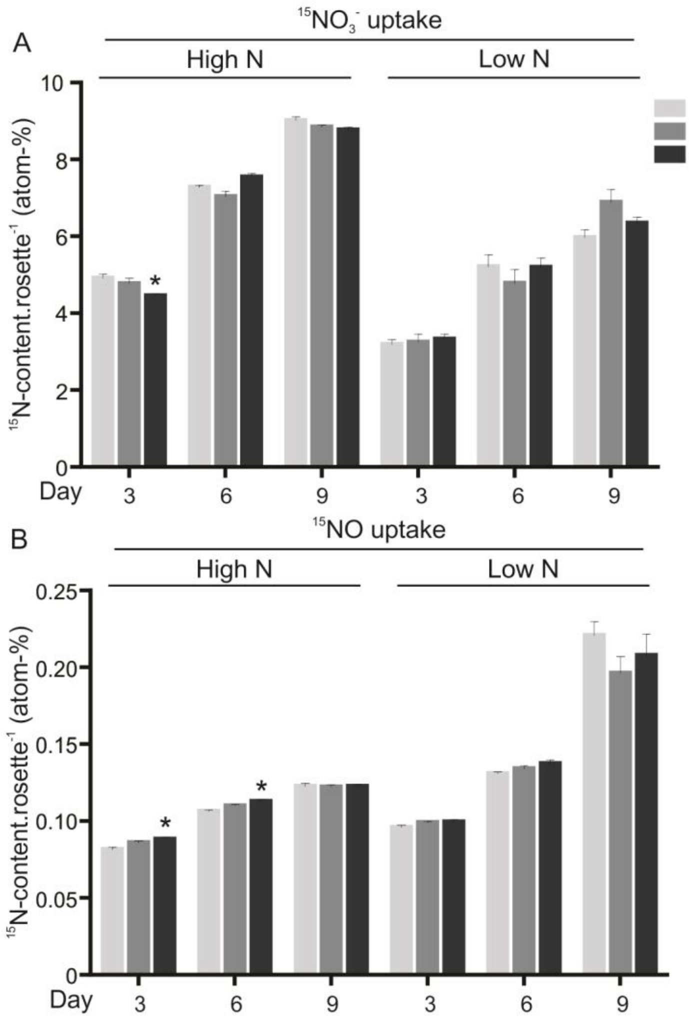


**Figure S2: Kinetics of $^{15}NO_3^-$ and atmospheric $^{15}NO$ uptake in SyNOS-transgenic plants**

Plants were grown in high and low N conditions as described in Fig. 4. Kinetics of $^{15}NO_3^-$ uptake (A) and atmospheric $^{15}NO$ incorporation (B) at 3, 6 and 9 days after switching N solutions. Values are means ± SE (n=6). Asterisks indicate significant differences in SyNOS lines compared to wt in the same N condition (ANOVA, Dunnett's post hoc test, *$p<0.05$).

**Table S1: Primers used for PCR analysis**

**PCR**

| Gene Name | Forward primer 5'- 3' | Reverse primer 5'- 3' |
|---|---|---|
| SyNOS | CGGATCCATGCTTGTCAACGACTC TCGTCCTACCGTAGAAGCGCACGT TCTTTCTGTT | GCTGCAGTCACAGGTCCTCCTCTG AGATCAGCTTCTGCTCCAAGTTGG CTAGCCATTT |
| Actin | AATCTCCGGCGACTTGACAG | AAACCCTCGTAGATTGGCACA |

**Real Time PCR**

| Gene Name | Forward primer 5'- 3' | Reverse primer 5'- 3' |
|---|---|---|
| SyNOS | TCGGCAGCACCGTCTATGAA | GTGCTGTCGGCTCCGAGAAT |
| NiA1 | GGTTCGAGCCTGGGACGAGT | TTGCCATCCACCCACCCGAC |
| NiR | AGATCCGTCCTTGTCGCCGC | TGGCTCGAGACGATCTGCGT |
| GS2 | CGCTTCGCCACAAGGAGCAC | TTCGCCTCGGTGTCACGTCC |
| GOGAT | CCGCCTCCTCACCACGACAT | GACTAGCCCCGGTTCCACCA |
| Actin | GCCATCCAAGCTGTTCTCTC | GAAACCCTCGTAGATTGGCA |
